# Wnt/β-catenin signaling improves oxidative metabolism in skeletal muscle of obese ob/ob mice

**DOI:** 10.1101/2024.01.23.576856

**Authors:** Eleni Christodoulou-Vafeiadou, Céline Lepeigneux, Aurore Naviere, Julien Pujol, Fadila Benhamed, Pascal Maire, Isabelle Guillet-Deniau

**Affiliations:** Institut Cochin, INSERM U1016, CNRS UMR 8104, Université Paris Cité. Paris. France

**Keywords:** : SREBP-1c, Wnt/β-catenin signaling, muscle, oxidative metabolism, obesity, diabetes

## Abstract

**Background:** Canonical Wnt signaling is involved in many physiological and pathological states. As it regulates lipid metabolism and glucose homeostasis, its misregulation may lead to the development of diabetes and obesity. We have already reported that activation of the Wnt/β-catenin canonical signaling pathway increased insulin sensitivity and prevented lipid deposits in rat skeletal muscle through a reciprocal regulation of Wnt10b and the lipogenic factor SREBP-1c.

**Results:** Here we have studied the role of Wnt/β-catenin canonical signaling in skeletal muscle of genetically obese and diabetic (ob/ob) mice and their control ob/+ mice. We showed that *Wnt10b* and *SREBP-1c* expressions were conversely regulated in cultured mouse myoblasts isolated from lean ob/+ or obese ob/ob mice. Activation of the Wnt/β-catenin pathway using Wnt10b overexpression or the selective GSK3 inhibitor 6-Bromo-indirubin-3’oxime (BIO) was sufficient to decrease lipogenic genes expression in cultured myoblasts isolated from control and obese mice. In vivo, we performed direct electrotransfection of Wnt10b cDNA or BIO injections in *Tibialis Anterior* (TA) muscles of ob/ob and ob/+ mice. Both up-regulated *Wnt10b* gene expression and down-regulated *SREBP-1c* expression. Canonical Wnt signaling increased slow Myosin Heavy Chain-I (MHC-I) oxidative fiber number as well as fast Myosin Heavy Chain-IIA (MHC-IIA) oxidative fiber number, while decreasing fast glycolytic fiber number in TA muscle. In addition, Wnt signaling increased mitochondrial oxidative metabolism and respiratory reserve capacity by 2- and 3-fold in myotubes cultured from ob/ob and ob/+ mice muscles respectively. Surprisingly, the activation of the Wnt pathway was sufficient to reduce hyperglycemia by 30% within 3 weeks in ob/ob mice.

**Conclusions:** Our results show that activation of Wnt/β-catenin signaling in skeletal muscle induced a shift towards a more oxidative metabolism in myofibers, thus mimicking the effects of exercise training. Wnt10b could be a valuable candidate to develop therapeutic drugs for the treatment of obesity and/or type 2 diabetes.

## BACKGROUND

### The Wnt/β-catenin pathway

The Wnt pathway is essential for embryonic development, tissue integrity, and stem cell activity [1, 2] but alterations in Wnt signaling can lead to the genesis of cancer [3]. While the 19 different Wnt ligands expressed in both human and mouse can activate several canonical and non canonical Wnt pathways, Wnt10b specifically activates canonical Wnt/β-catenin signaling and thus triggers β-catenin/LEF/TCF-mediated transcriptional programs in the immune system, adipose tissue, mammary gland, bone and skin, where it controls stemness, pluripotency and cell fate. Dysregulation of these processes causes breast cancer, osteoporosis, but also metabolic disorders such as type 2 diabetes and obesity [4, 5].

### Link between Wnt signaling and obesity/type 2 diabetes

The first indication of a possible link between Wnt signaling and metabolic diseases came from studies that reported a C256Y mutation in WNT10b gene associated with obesity [4], and a single nucleotide polymorphism in the WNT5b gene in a Japanese cohort that developed type 2 diabetes [6]. In humans, another study pointed out the association between variants of the Transcription factor 7-like 2 (TCFL2 or TCF4) and the pathogenesis of type 2 diabetes [5]. In vitro, a link between cellular glucose sensing and Wnt/β-catenin signaling was reported in macrophages [7], indicating that this pathway could be inappropriately activated in diabetic hyperglycemic or obese subjects. In a rat model of high-fat-diet-induced obesity, injection in *Tibialis Anterior* and *Soleus* muscles of a recombinant adeno-associated virus (rAAV) vector encoding murine Wnt10b cDNA reduced global and muscular fat deposits and body weight. In addition, the overexpression of Wnt10b was accompanied by reduced hyperinsulinemia, triglycerides, leptin and adiponectin plasma levels as well as improved glucose homeostasis [8]. On the other hand, transgenic mice expressing Wnt10b under the FABP4 promoter have less adipose tissue and are resistant to diet-induced obesity [9]. In addition, these mice do not gain significant body weight on the ob/ob background. FABP4-Wnt10b mice are more glucose tolerant and insulin sensitive than control mice on the lethal yellow agouti background [10]. Similarly, it was suggested that diet-induced obesity in C57Bl6/J mice was correlated to the up-regulation of SFRP5, Dapper, Dickkopf and Naked1 in adipose tissue, four genes known for their inhibitory effects on Wnt signaling [11].

### Role of Wnt signaling in the balance adipogenesis/myogenesis

All these data strongly suggest that Wnt signaling could be involved in the regulation of glucose homeostasis, particularly in insulin sensitive tissues such as skeletal muscle. Although it has been shown that Wnt10b represses adipogenesis [12] and activates myogenesis [13, 14] less is known about the involvement of Wnt10b in the etiology of metabolic disorders. Several studies indicate a strong correlation between accumulation of intramyocellular lipids and insulin sensitivity in humans, not only in diabetic patients but also in glucose-tolerant and -intolerant individuals with or without obesity [15, 16]. In addition, muscle-specific inhibition of Armadillo, the *Drosophila* β-catenin ortholog and transcriptional effector of Wg signaling, evoked an obese phenotype in flies [17].

We have previously shown that increased intramuscular lipid accumulation associated with muscle insulin resistance in obesity or type 2 diabetes arose partly from de novo fatty acid synthesis in rat skeletal muscle [18]. We also reported that activation of the Wnt/β-catenin canonical pathway increased insulin sensitivity in rat muscle cells and prevented lipid deposits through a reciprocal regulation of Wnt10b and SREBP-1c [19], a lipogenic factor that mediates insulin’s effects on hepatic [20, 21] and skeletal muscle gene expression in humans [22] and rodents [23]. Furthermore insulin resistance in obese and diabetic patients is strongly correlated with the activity of glycolytic or oxidative enzymes in skeletal muscle [24], and muscle oxidative capacity as well as insulin sensitivity are decreased in relation with a switch from slow oxidative fibers to fast glycolytic fibers in elderly or sedentary subjects [25, 26]. In addition, Wnt signaling modulates both the number of terminally differentiated myogenic cells and the intricate slow/fast patterning of the avian [27] and mouse limb musculature [28].

To pinpoint the role of Wnt/β-catenin signaling in muscle metabolism, we have activated the Wnt/β-catenin pathway in skeletal muscles of genetically obese and diabetic ob/ob mice, as well as in primary cultures of satellite cells isolated from obese mice muscles. Here we show for the first time that the activation of the Wnt/β-catenin pathway through Wnt10b overexpression or treatment with 6-Bromoindirubin-3’ Oxime (BIO), a specific inhibitor of GSK-3, is sufficient to increase slow oxidative fiber number in TA muscle and to decrease hyperglycemia in ob/ob mice. The switch in fiber type towards oxidative fibers is accompanied by an increase in the mitochondrial oxidative metabolism as well as in the respiratory reserve capacity.

Our results strongly suggest that Wnt10b could be a valuable candidate to develop therapeutic drugs for the treatment of obesity and/or type 2 diabetes.

## METHODS

### Ethics Statement

In vivo experimentation was carried out in strict accordance with the Europeen convention STE 123 and the French national charter on the Ethics of Animal Experimentation. Protocols were approved by the Ethical Committee of Animal Experiments of the Institut Cochin, CNRS UMR 8104, INSERM U1016. Surgery was performed under ketamine/xylazine anesthesia, all efforts were made to minimize suffering.

### Animals

Six-week-old male C57BL6J/ob/+ and C57BL6J/ob/ob mice (Janvier, France) were maintained in the animal room of Institut Cochin (agreement n° B 75-14-02). They received a standard laboratory chow and water ad libitum until sacrifice. Before in vivo electrotransfection of *Tibialis Anterior* (TA) muscles, mice were anesthetized with a ketamine/xylazine cocktail (90 mg/kg Ketamine, 4.5 mg/kg Xylazine). Three weeks later, animals were killed by cervical dislocation and TA and *Soleus* (Sol) muscles were harvested immediately after sacrifice.

### In vivo activation of Wnt/β-catenin signaling

To determine the effects of Wnt/**β**-catenin signaling on skeletal muscle in vivo, we activated this pathway in *Tibialis Anterior* (TA) of ob/+ and ob/ob mice using in vivo electrotransfection (IVE) of a plasmid encoding a mouse Wnt10b cDNA, or through injections of BIO.

#### In Vivo Electrotransfection

A mix of 4 μg of pcDNA3.1 plasmid containing the mouse Wnt10b cDNA (provided by Dr Bernardi, INRA, Montpellier) and 1 μg of pAcGFP-Nuc that encodes a nuclear GFP (Clontech) in 0.9% NaCl was electrotransfected directly into TA muscles of anesthetized ob/+ and ob/ob mice according to the protocol described by Grifone et al.[29]. The contralateral TA was considered as a control and electrotransfected with 4 μg of pAcGFP-Nuc. Three weeks later, mice were sacrificed and blood and tissue samples were harvested.

#### BIO Injections

BIO (20 nmoles, (Calbiochem®) was directly injected into the TA muscle of ob/+ and ob/ob mice under a brief fluothane anesthesia, while the same amount of Me-BIO, an inactive form of BIO [30], was injected into the contralateral TA muscle. Injections were performed twice a week for 3 weeks. For systemic administration, six week-old mice received intraperitoneal (IP) injections of BIO or Me-BIO (50 nmoles) twice a week for 7 weeks. Body weight and glycemia were measured every other day in both models.

### Primary culture of mouse muscle satellite cells

Three different cultures were performed from three different mice of each group (ob/+ and ob/ob mice). Satellite cells from mice hindlimb muscles were isolated and cultured in a controlled humidified atmosphere (7% CO_2_, 37 °C) as previously described [31]. Cells were fed with fresh medium the day after plating and every other day. At day 6, the medium was replaced by a differentiation medium (2% horse serum) and myoblasts differentiated into spontaneously contracting myotubes within 3 days, then serum was totally removed.

### Transfection of myotubes

Mouse myotubes were cultured as described above and antibiotics were removed the day before transfection. Cells were transfected with the recombinant pcDNA3.1 plasmid (6 μg/flask) containing mouse *Wnt10b* cDNA using Lipofectamine 2000 (Invitrogen) according to the manufacturer’s instructions. Efficiency of transcription was controlled using a Nikon epifluorescence inverted microscope (Nikon ECLIPSE TS100-F). Experiments were performed 48 h later.

### Cellular oxygen consumption studies

Myotubes cultured under physiological (5 mM, G5) or supra-physiological (25 mM, G25) glucose concentration were treated with 100 μM BIO for 48 h or transfected with Wnt10b cDNA. Mouse myotubes were trypsinized and resuspended in 3 ml of DMEM medium. Cell respiration was measured using a polarographic oxygen sensor in a 2-ml glass chamber (Oroboros Oxygraph 2K, Oroboros Instruments, Austria) as described in Gnaiger [32]. Basal respiration rate was determined at 37°C after stabilization of both temperature and O_2_ saturation. Oligomycin (0,5 μg/ml) was then added to inhibit ATPase and down-regulate the respiratory rate. Increasing concentrations (5 to 10 μM) of carbonyl cyanide m-chlorophenyl hydrazone (CCCP) was added to estimate the maximum (uncoupled) respiratory rate. Finally 1 μM potassium cyanide (KCN) was added to inhibit completely cellular respiration. Protein content was measured in cells and the respiratory rate was expressed as O_2_ flux (pmoles/second/μg protein). This technique allows to determine the mitochondrial respiratory reserve capacity (defined as maximal respiration speed minus basal respiration speed).

### PCR and Q-PCR

Total RNA was extracted using the TRIzol reagent (Invitrogen) according to the manufacturer’s instructions. First-strand cDNA was synthesized from 600 ng of total RNA in the presence of 100 U of SuperScript^TM^ II Reverse Transcriptase (Invitrogen). Semi-quantitative PCR was performed from cDNAs with Dream Taq (Fermentas) or Advantage® (Clontech) polymerase for 28–32 cycles. All primers were designed according to sequences in GenBank and β-actin was used as an internal control. For quantitative PCR, cDNAs were amplified using the LightCycler® 480 SYBR Green I Master (Roche). The relative mRNA abundance was calculated using the LightCycler® software and normalized to Cyclophillin and Tata Binding Protein (TBP) mRNAs. Results were obtained from four to five independent RNA samples from individual experiments, each tested in duplicate.

Oligonucleotide primers relative to the genes of interest are listed in Additionnal file 1.

### Western Blot analysis

Proteins (30 μg) were separated by SDS-PAGE in 9,5% gels and electrotransferred onto nitrocellulose. Membranes were blotted with the relevant antibody and then revealed using the ECL system (Supersignal, Pierce). Signals were scanned and quantified using ImageJ software.

### Antibodies

SREBP-1c was detected using a monoclonal antibody raised against the N-terminus of human SREBP-1 (NeoMarkers). Anti-FAS was a gift from Dr I. Dugail (Unité INSERM 465, Paris, France). Rabbit monoclonal antibody against β-catenin was from Epitomics. Monoclonal antibody against active β-catenin was from Chemicon. Antibodies raised against β-tubulin and Wnt10b were from Santa-Cruz Biotechnology. Mitochondrial respiratory chain subunit B6 of Complex I and core 1 of Complex III antibodies were gifts from Dr. Anne Lombes, INSERM U1016, Paris, France. Anti-porin was purchased from the Calbiochem company. For immunohistochemistry, anti-laminin and anti-MHC-I antibodies were purchased from SIGMA. Anti-MHC-IIA was a kind gift of Dr Schiaffino (Department of Biomedical Sciences, University of Padova,Italy).

### Histological and immunohistochemical analysis

TA muscles were harvested and immediately frozen after sacrifice. Serial muscle cryosections (7 μm thick) were made in the median part of the TA muscle. For fiber typing, sections were fixed with 2% paraformaldehyde (PFA) and saturated in a blocking solution (horse serum - Vector Laboratories). Sections were incubated for 2h30 with primary antibodies (anti-laminin, 1/250; anti-MHC-I, 1/1000; anti-MHC-IIA, 1/10). Image acquisition was performed using a Nikon inverted microscope (Nikon ECLIPSE TS100-F) equipped with a Nikon camera head DS-5M. Stained slices were observed under a epifluorescence macroscope (Nikon AZ 100M) connected to a Nikon DS-Ri1 high-resolution color camera. Images were analyzed using the imaging software NIS-Elements BR 3.00. Total fiber number of TA serial sections from 5 ob/+ and 6 ob/ob mice were counted using ImageJ software. Fiber cross-sectional area (CSA) was determined using MetaMorph software (Molecular Devices) after immunohistochemical labeling of cryostat sections with anti-laminin antibody. At least 8 to 10 slices from three different animals were analyzed for each group.

### Intramyocellular Lipid Accumulation

Intramyocellular lipids were detected in myotubes using Oil Red O staining according to Koopman et al [33]. Nuclei were counterstained with DAPI (Molecular Probes) then fluorescence was observed using a Nikon TS100 fluorescence microscope.

### Statistical Analysis

Results are expressed as means ± S.E. Experimental conditions were compared to control conditions using ANOVA tests with GraphPad Prism 5 software for Mac Os X. Differences were considered significant for p*<*0.05.

## RESULTS

### ACTIVATION OF WNT/β-CATENIN SIGNALING IN MOUSE MYOTUBES

#### Characterization of myoblasts isolated from ob/+ and ob/ob mice muscles

Myoblasts were allowed to differentiate into contracting myotubes within 10 days. Myosin heavy chain genes were differentially expressed : during differentiation, ob/+ myoblasts expressed mainly *Myh7* gene that encodes the slow oxidative isoform MHC-I, whereas ob/ob myoblasts expressed also *Myh4* encoding the fast glycolytic isoform MHC-IIb (Fig. 1A, a, b). This result suggests that obesity and type 2 diabetes could induce profound modifications in muscle fiber metabolism in the ob/ob mice. Similarly to what we observed in wild type murin myoblasts [19], *Wnt10b* and *Srebp-1c* genes had inverse expression patterns in myoblasts isolated from ob/+ and ob/ob mice. *Wnt10b* was only expressed in myoblasts, whereas *Srebp-1c* expression was only detected in myotubes (Fig. 1A, c).

**Fig. 1.**
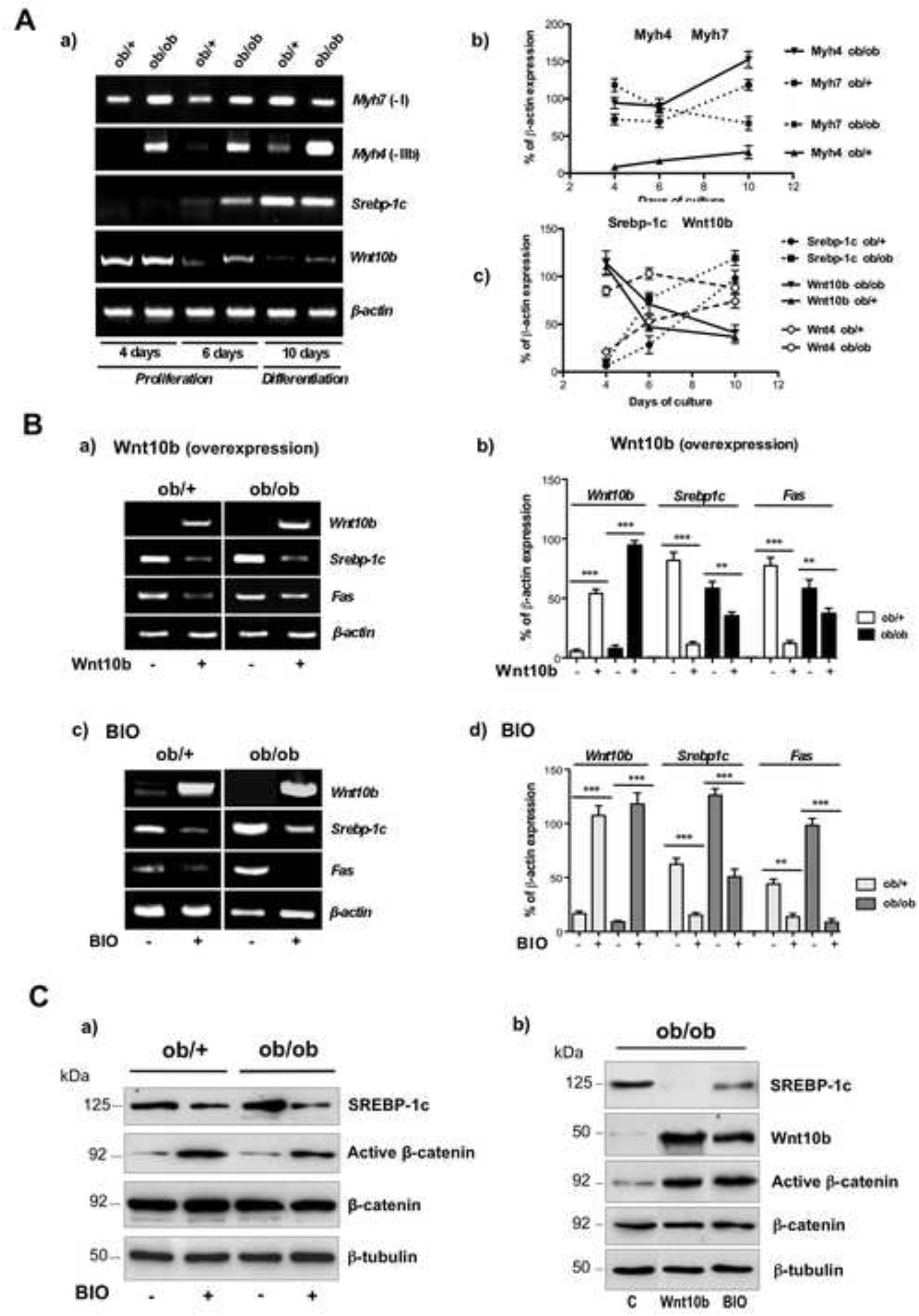
(A) Relative mRNA expression of Wnt10b and SREBP-1c during differentiation of cultured satellite cells from ob/ob mice. Semi-quantitative PCR was performed on satellite cells allowed to proliferate for 6 days then to differentiate into myotubes until day-10. ob/+ myotubes expressed *Myh7* (encoding the MyHC-I slow isoform) whereas ob/ob myotubes expressed *Myh4* (encoding the fast MyHC-IIb isoform) **(a, b)**. *Wnt10b* and *Srebp-1c* showed inverse expression profiles during differentiation of both ob/+ and ob/ob satellite cells **(a, c)**. **(B)** *Wnt10b* overexpression was sufficient to down-regulate *Srebp-1c* and *Fas* mRNA expression by 5-fold and 2-fold in ob/+ and ob/ob mice myotubes respectively **(a, b)**. BIO induced a 5-fold increase in *Wnt10b* mRNA expression that led to a 2-to 3-fold decrease in *Srebp-1c* and *Fas* mRNA expression **(c, d)**. **(C)** Western blot showing activation of β-catenin 24h after addition of 100 μM BIO in ob/+ and ob/ob mice myotubes (a). Transfection of Wnt10b or addition of BIO activated β-catenin in ob/ob mice myotubes (b). Blots are representative of three independent experiments.

#### Wnt/β-catenin signaling decreased lipogenic genes expression in mouse myotubes

Myotubes cultured from ob/+ and ob/ob mice muscles were transfected with a plasmid encoding mouse *Wnt10b* cDNA or treated with 100 μM BIO as described in methods. RNA was isolated 48 h later. *Wnt10b* overexpression through cDNA transfection or BIO addition was sufficient to drastically decrease the expression of the lipogenic genes *Srebp-1c* and *Fatty acid synthase* (*Fas*) in cells isolated from ob/+ and ob/ob mice (Fig.1B, a, b). Surprisingly, addition of BIO induced a 5-fold increase in *Wnt10b* mRNA expression that was concomitant with a 3-fold decrease in *Srebp-1c* expression in myotubes isolated from ob/+ and ob/ob mice (Fig.1B, c, d). These results confirm that *Wnt10b* and *Srebp-1c* are inversely expressed in cultured muscle satellite cells isolated from control mice as well as diabetic obese mice.

In order to determine whether the Wnt/β-catenin pathway was activated by Wnt10b and BIO in myotubes isolated from ob/+ and ob/ob mice, we checked the activation of β-catenin using an antibody raised against its active form. Wnt10b and BIO up-regulated active β-catenin protein level by 5-fold while SREBP-1c protein was down-regulated (Fig.1C, a, b). Taken together, these results prompted us to investigate the role of Wnt signaling in skeletal muscle of ob/ob mice in vivo.

### IN VIVO ACTIVATION OF WNT/β-CATENIN SIGNALING

#### Effect of Wnt signaling on mice body weight and glycemia

##### Activation of Wnt/β-catenin signaling in TA muscle

Three weeks after In Vivo Electrotransfection (IVE) of Wnt10b or BIO injections in the TA, no significant effect was observed on body weight of ob/+ or ob/ob mice (Fig. 2A, a). In contrast, BIO injections or Wnt10b IVE in the TA were sufficient to decrease hyperglycemia by respectively 27% and 29% in ob/ob mice, while glycemia of ob/+ mice was unchanged (Fig. 2A, b, c).

**Fig. 2.**
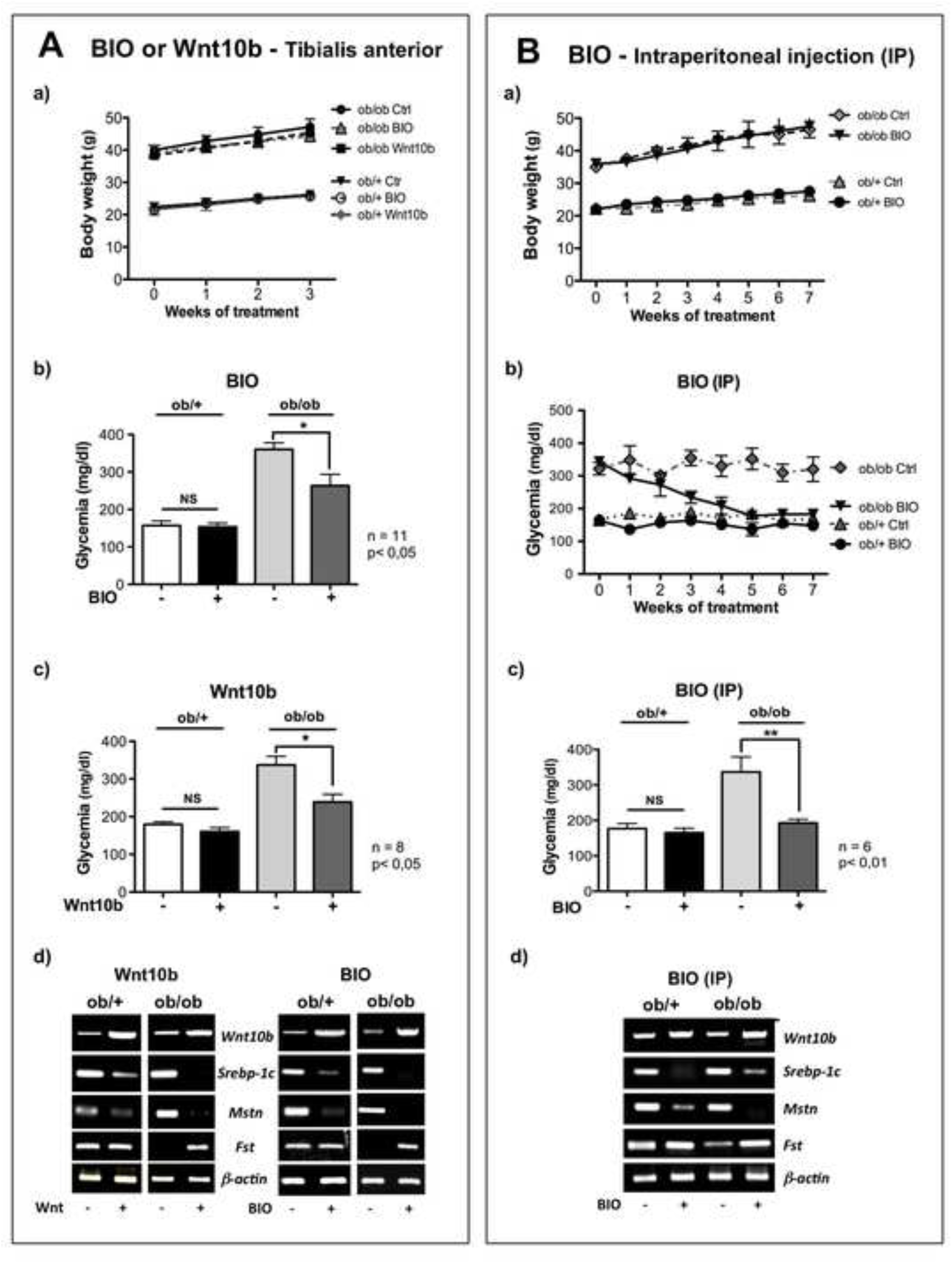
Effect of a local (A) or systemic (B) activation of Wnt signaling in ob/ob mice. **(A) Local activation of Wnt signaling.** The TA muscle from the right hindlimb was electrotransfected with Wnt10b cDNA or injected with 20 nmoles BIO twice a week for 3 weeks, while the contralateral TA muscle was injected with PBS or Me-BIO as a control. **(a)** Activation of Wnt signaling had no effect on body weight of ob/+ and ob/ob mice. In contrast, BIO injection **(b**) or Wnt10b electrotransfection **(c)** was sufficient to decrease glycemia of ob/ob mice by 29% and 27% respectively, whereas it has no effect on the glycemia of ob/+ mice. **(d)** *Wnt10b* overexpression or BIO injection down-regulated mRNA expression of the lipogenic gene *Srebp-1c* as well as the expression of Myostatin *(Mstn),* whereas the expression of Follistatin *(Fst)* was increased in the TA muscle of ob/ob mice. **(B) Systemic activation of Wnt signaling.** Mice received intraperitoneal (IP) injections of 50 nmoles BIO twice a week for 7 weeks. **(a)** BIO had no effect on ob/+ and ob/ob mice body weight. **(b, c)** BIO decreased glycemia of ob/ob mice by 40% whereas glycemia of ob/+ mice remained unchanged. **(d)** Systemic injection of BIO was sufficient to up-regulate *Wnt10b* mRNA expression in TA muscles of ob/+ and ob/ob mice, while *Srebp-1c* and *Mstn* mRNA levels were down-regulated. In contrast, *Fst* expression was increased only in the TA muscle of ob/ob mice.

##### Systemic activation of Wnt/β-catenin signaling

In order to study the effect of a systemic activation of Wnt signaling on body weight and glycemia, BIO was injected intraperitoneally (IP) to the mice twice a week for 7 weeks. Systemic administration of BIO had no effect on body weight of ob/+ and ob/ob mice (Fig. 2B, a). In contrast, BIO decreased by 40% hyperglycemia of ob/ob mice within 5 weeks of treatment, while glycemia of ob/+ mice was unaffected (Fig. 2B, b, c). These results show that local (TA muscle) or systemic (IP injection) activation of Wnt/β-catenin signaling was able to decrease glycemia of obese hyperglycemic ob/ob mice by mechanisms that remain to be determined. In contrast, glycemia of normoglycemic ob/+ mice was unchanged.

#### Local effects of Wnt signaling

As Wnt proteins are known to have autocrine and paracrine effects [34] we checked whether activation of Wnt signaling could act not only on TA itself, but also on neighboring muscles.

##### Effect on muscle weight

TA and *Soleus* (Sol) muscles were weighted 3 weeks after Wnt10b IVE or BIO injections. Wnt10b overexpression or BIO injections did not significantly affect the weight of TA and Sol muscles in ob/+ as well as in ob/ob mice. Similarly, IP injections of BIO had no effect on TA and Sol weight (data not shown).

##### Effect on gene expression

Three weeks after Wnt10b IVE or BIO injections in TA of ob/+ and ob/ob mice, mRNA expression was assessed using semi-quantitative RT-PCR. As expected, Wnt10b transfection and BIO injections decreased *Srebp-1c* gene expression in TA muscles of both ob/+ and ob/ob mice, but also down-regulated Myostatin *(Mstn)* expression (a TGF-β family member that inhibits muscle differentiation and growth [35] (Fig 2A, d). Activation of Wnt signaling also up-regulated Follistatin *(Fst)* (a factor that binds and neutralizes members of the TGF-® superfamily such as Activin and Myostatin [36]) only in the TA of ob/ob mice (Fig. 2A, d). As it was the case in cultured myotubes isolated from ob/+ and ob/ob mice (Fig. 1B, c), IP injections of BIO increased *Wnt10b* expression in TA muscle concomitantly with the down-regulation of *Srebp-1c* and *Mstn* expression. Furthermore, *Fst* gene expression was up-regulated in TA muscle of ob/ob mice (Fig. 2B, d).

Taken together these results demonstrate that Wnt10b and BIO had similar effects on glycemia and lipogenic genes expression in ob/ob mice. As BIO induced the up-regulation of *Wnt10b* gene expression in TA muscle, all further experiments were performed after IVE of *Wnt10b* cDNA.

#### Effect of Wnt signaling on muscle plasticity

##### Determination of fiber size, number and type distribution

Three weeks after Wnt10b IVE, TA from ob/+ and ob/ob mice were sliced and fiber size was evaluated using cross section area (CSA) measurement after staining of the basal lamina with an antibody raised against Laminin. Total fiber number was significantly lower in the TA of ob/+ mice (1265 ± 59) (Fig. 3A, a) than in the TA of ob/ob mice (1610 ± 197) (Fig. 3A, b). Wnt10b IVE did not affect total fiber number neither in ob/+ mice (1203 ± 131) (Fig. 3A, a) nor in ob/ob mice (1697 ± 258) (Fig. 3A-b). Fiber size of ob/+ TA muscle ranged from 30 to 1200 μm^2^ (Fig. 3A, c) while the TA of ob/ob mice was mainly constituted of small fibers (60% of 101-300 μm^2^) (Fig. 3A, d). Wnt10b overexpression increased by 2.5 fold small fiber number (30-100 μm^2^) in the TA of ob/ob mice, while the number of fibers ranging from 101-500 μm^2^ was significantly decreased (Fig 3A, d). In contrast, Wnt10b overexpression had no effect on fiber size in the TA of ob/+ mice (Fig 3B-c).

**Fig. 3.**
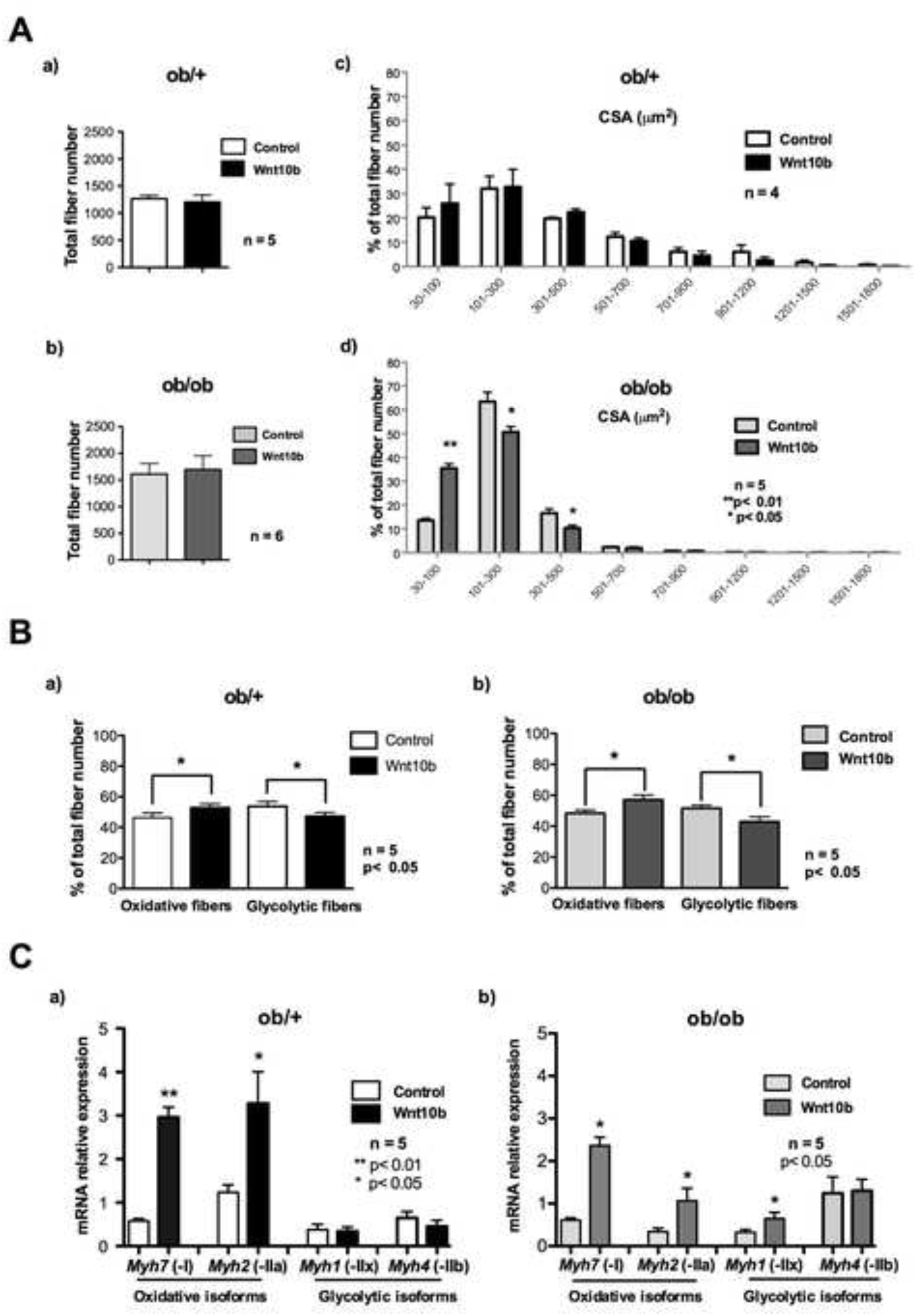
Wnt10b overexpression increased oxidative fiber number in the TA muscle. **(A)** Wnt10b overexpression did not change total fiber number in the TA muscle of ob/+ **(a)** and ob/ob **(b)** mice. CSA measurements were performed using MetaMorph software on TA slices stained with Laminin three weeks after in vivo electrotransfection. **(c)** Wnt10b overexpression did not affect the distribution of fibers in the TA of ob/+ mice. **(d)** On the contrary small fiber number was increased by 2.5 fold (30-100μm^2^) in the TA of ob/ob mice. **(B)** Oxidative (SDH positive) fiber number was increased by 6,5% and 8,5% in ob/+ **(a)** and ob/ob **(b)** mice respectively, whereas glycolytic fiber number was decreased of the same amount. **(C)** Determination of myosin mRNA expression by quantitative PCR. Wnt10b overexpression increased mRNA of oxidative *Myh7* (-I) and *Myh2* (-IIA) isoforms by 4-fold and 2,5-fold respectively in the TA of ob/+ **(a)** and ob/ob **(b)** mice. In contrast, mRNA encoding glycolytic *Myh1*(-IIX) and *Myh4 (*-IIB) isoforms was unaffected **(a), (b)**.

##### Determination of the fiber-specific glycolytic-to-oxidative ratio

Three weeks after Wnt10b IVE, TA from ob/+ and ob/ob mice were sliced and sections were stained for mitochondrial Succinate DeHydrogenase activity (SDH) which reflects the oxidative capacity of muscle fibers (Fig. 4A). Wnt10b overexpression increased oxidative fiber number by 6,5% and 8,7% in the TA of ob/+ (Fig. 3B, a) and ob/ob mice respectively (Fig. 3B, b), whereas SDH negative fiber number was decreased of the same amount. These results suggest that activation of Wnt/β-catenin signaling induced a fiber switch towards a more oxidative metabolism in TA muscles.

**Fig. 4.**
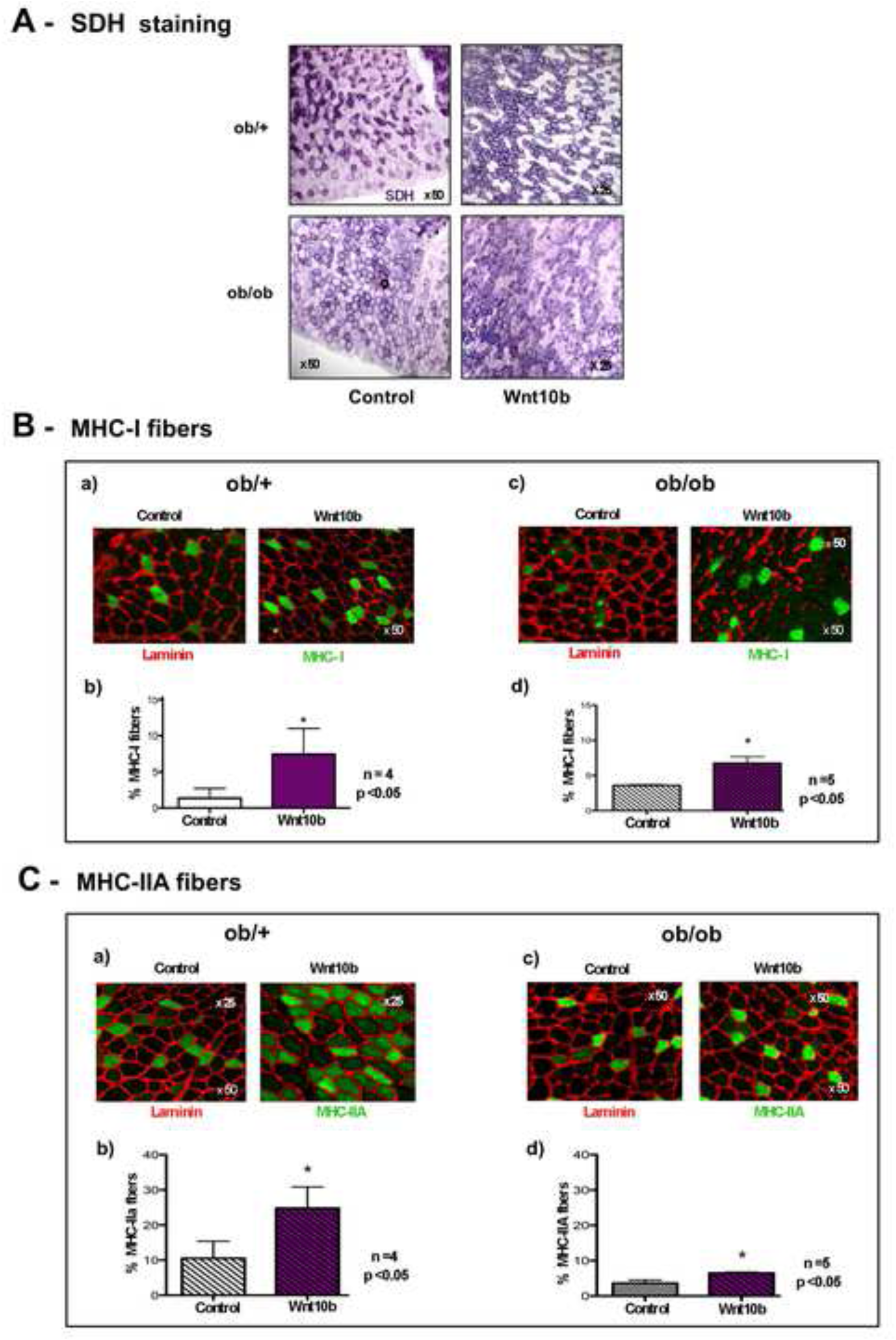
Quantification of oxidative MHC isoforms 3 weeks after Wnt10b electrotransfection. **(A)** Succinate dehydrogenase (SDH) assay showing a stronger staining (purple) in the more oxidative fibers after Wnt10b electrotransfection in the TA muscle of ob/+ and ob/ob mice. **(B)** Wnt10b overexpression increased oxidative MHC-I fiber number by 4-fold and 2-fold in the TA of ob/+ **(a), (b)** and ob/ob **(c), (d)** mice respectively. **(C)** In addition oxidative MHC-IIA fiber number was increased by 2-fold in ob/+ **(a), (b)** and ob/ob **(c), (d)** mice. MHC-I and MHC–IIA oxidative fibers were stained in green. Laminin was counterstained in red. Quantification of fiber type was performed using ImageJ software.

##### Q-PCR quantification of mRNA encoding MHC isoforms

Muscle fibers express different Myosin Heavy Chains (MHC) isoforms depending on their metabolism. Slow oxidative fibers express mainly the MHC-I isoform (which is encoded by the *Myh7* gene), while fast oxidative fibers express preferentially the MHC-IIA isoform (encoded by the *Myh2* gene). Wnt10b IVE increased oxidative *Myh2* (-IIA) mRNA expression by 3-fold and 2.5-fold respectively in the TA of lean (Fig.3C a) and obese mice (Fig. 3C, b). It is noteworthy that among glycolytic isoforms, *Myh1* mRNA (-IIX) was also increased by 2-fold in the TA of ob/ob mice, whereas *Myh4* (-IIB) was unchanged in both mice. In addition, *Myh7* was enhanced by 4-fold and 3-fold respectively by Wnt10b IVE in the TA of ob/+ and ob/ob mice (Fig. 3C, a, b). Taken together, these findings confirm that Wnt/β-catenin signaling not only induced a more oxidative metabolism, but also modulated *Myh* gene expression particularly in the TA of obese mice.

##### Expression of MHC isoforms upon activation of Wnt signaling

To confirm a fiber-type switch, TA sections were stained with antibodies raised against slow MHC-I or fast MHC-IIA isoforms and fibers positive for each antibody were counted. Wnt10b overexpression increased MHC-I positive fibers by 4-fold and 2-fold respectively in the TA of ob/+ (Fig. 4B, a, b) and ob/ob mice (Fig. 4B, c, d) while the number of MHC-IIA positive fibers was enhanced by about 2-fold both in the TA of lean and obese mice (Fig. 4C).

### THE WNT/β-CATENIN PATHWAY INCREASED MITOCHONDRIAL OXIDATIVE METABOLISM IN SKELETAL MUSCLE

#### Expression of factors involved in mitochondrial metabolism

We wonder whether the increase in oxidative fiber number observed after activation of Wnt signaling could be related to a change in mitochondrial metabolism. To assess this hypothesis, we measured the expression of *Pgc-1α* and its targets *Hif-2α* and *Mef-2c* (two factors involved in the switch of fibers towards an oxidative phenotype [37]) as well as *Pgc-1β* after activation of Wnt signaling through the overexpression of Wnt10b in TA muscles.

##### Q-PCR quantification of Pgc-1α, Pgc-1β, Hif-2α and Mef-2c

Total RNA was extracted to perform Q-PCR quantifications. Fig. 5-A shows that overexpression of Wnt10b induced a significant increase in *Pgc-1α* mRNAs in the TA of ob/ob mice (3.74 ± 1.12 versus 1.46 ± 0.18) as well as in the TA of ob/+ mice (5.05 ± 0.57 versus 3.74 ± 0.43) whereas *Pgc-1β* mRNA expression was unchanged. In the same manner, *Hif-2α* expression was increased in TA muscles of ob/+ (1.08 ± 0.09 versus 0.69 ± 0.08) and ob/ob mice (1.84 ± 0.40 versus 0.73 ± 0.12). In contrast, *Mef-2c* expression was increased by 3-fold in the TA of ob/ob mice (2.17 ± 0.37 versus 0.77 ± 0.10), but not in the TA of ob/+ mice (1.41 ± 0.42 versus 1.40 ± 0.46).

**Fig. 5.**
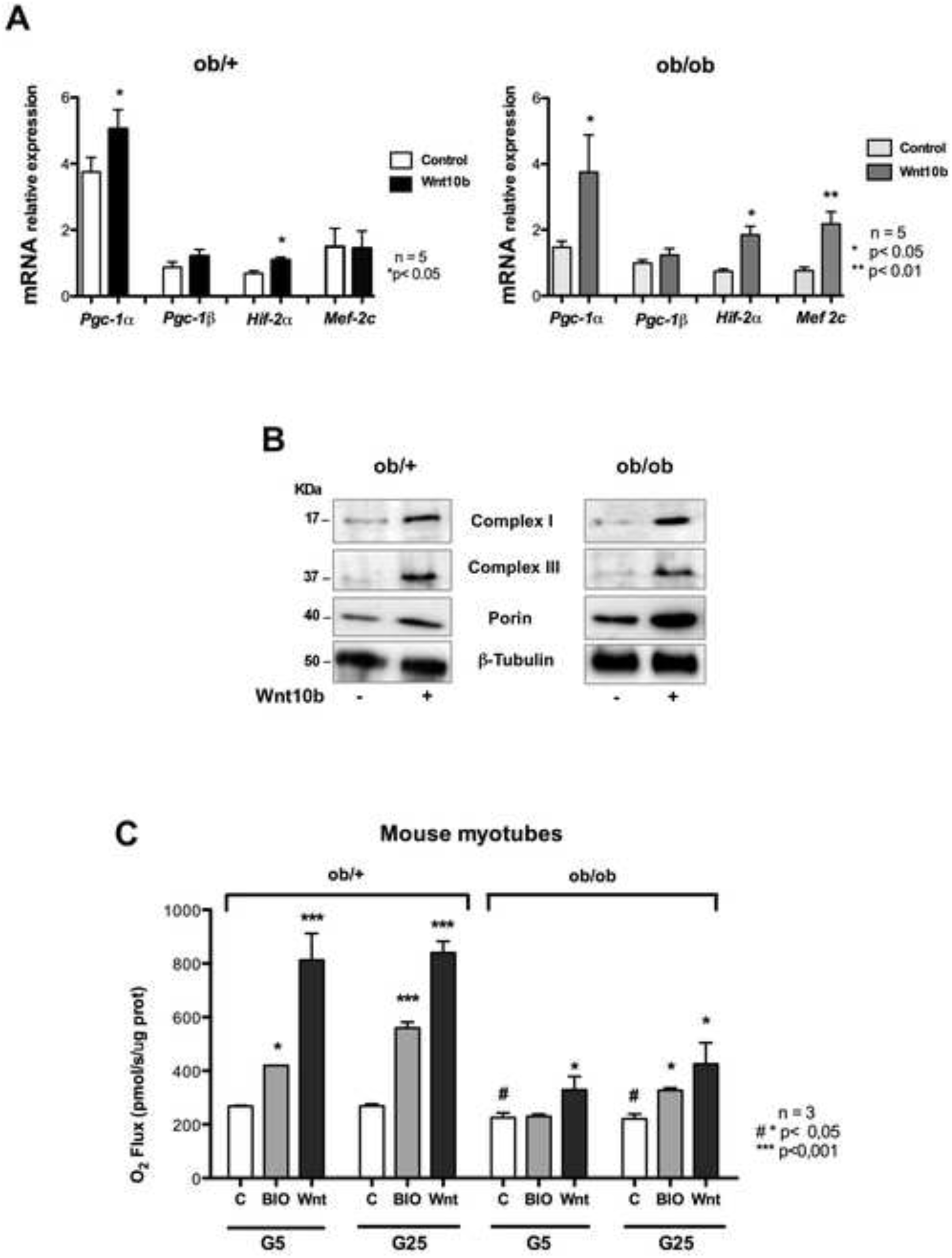
Effect of Wnt signaling on muscle oxidative metabolism. **(A)** Q-PCR quantification of *Pgc-1α*, *Pgc-1β*, *Hif**-**2α* and *Mef-2c* in the TA of ob/+ and ob/ob mice. *Pgc-1α* was increased by 30% in the TA of ob/+ mice and by 2-fold in the TA of ob/ob mice. In contrast, *Pgc-1β* was not affected neither in ob/+ or ob/ob mice. **(B)** Western Blot analysis showing that overexpression of Wnt10b in the TA up-regulated mitochondrial proteins belonging to the respiratory chain (B6 subunit of complex I and core 1 subunit of complex III) as well as porin (a protein of the outer membrane) in both ob/+ and ob/ob mice TA muscles. Western Blots are representatives of three independent experiments. **(C)** Effect of Wnt signaling on cellular respiration. Mitochondrial respiration was measured in mouse myotubes using the Oroboros Oxygraph 2K. Results are expressed as mitochondrial respiratory reserve capacity (maximal respiration speed minus basal respiration speed). Activation of Wnt signaling through BIO or Wnt10b increased by 2-to 3-fold the respiratory reserve capacity of ob/+ myotubes cultured in 5 mM (G5) or 25 mM (G25) glucose concentration. In contrast, the respiratory reserve capacity was increased by only 30% to 50% in myotubes of ob/ob mice cultured in G5 and G25, respectively. Results are means ± s.e. of 3 independent experiments performed in triplicate.

#### Wnt Signaling up-regulated mitochondrial proteins of the respiratory chain

In parallel, total protein was isolated and western blot was hybridized using antibodies raised against mitochondrial proteins belonging to complexes of the respiratory chain. Wnt10b overexpression enhanced the B6 subunit of Complex I, as well as the core-1 subunit of Complex III in the TA muscle of lean and obese mice. We also found that porin, a protein that allows small organic molecules transport through the outer membrane, was enhanced in both models (Fig. 5B). These results prompted us to evaluate the respiratory capacity of mitochondria under activation of Wnt signaling by measuring oxygen consumption of cultured myoblasts isolated from muscles of ob/+ and ob/ob mice.

#### Effect of Wnt signaling on cellular respiration

Mitochondrial respiratory control through the phosphorylation system is partially or fully released by pathophysiological uncoupling and dyscoupling. In order to determine whether Wnt signaling could affect cellular respiration, we performed high-resolution respirometry on mouse intact myotubes using the OROBOROS Oxygraph 2K as described in Methods.

##### Effect of Wnt signaling on the cellular respiration of mouse myotubes

The basal respiratory reserve capacity of myotubes isolated from ob/ob mice was slightly but significantly decreased as compared to ob/+ mice. Furthermore, activation of Wnt signaling through Wnt10b overexpression increased by 3-fold the respiratory reserve capacity of myotubes from ob/+ mice whatever the glucose concentration, whereas BIO addition induced a 2-fold increase (Fig. 5-C). The respiratory reserve capacity of myotubes isolated from ob/ob mice overexpressing Wnt10b was increased by 50% and 90% in 5 mM (G5) and 25 mM (G25) glucose concentrations respectively. In contrast BIO increased the basal respiratory reserve capacity only in myotubes cultured in high glucose concentration (G25) (Fig. 5-C). These results show that activation of Wnt signaling increased mitochondrial activity under physiological and pathophysiological states.

### ROLE OF WNT/β-CATENIN SIGNALING ON GLUCIDE AND LIPID METABOLISM

#### Effect of BIO on insulin sensitivity of myotubes

As ob/ob mice were reported to show a strong liver and muscle insulin resistance [38], we analyzed the role of the Wnt/β-catenin pathway on insulin sensitivity of myotubes isolated from ob/+ and ob/ob mice. Myotubes were treated with 100 μM BIO for 24h in the presence or absence of 10 nM insulin for the last 12h. Insulin increased SREBP-1c and FAS lipogenic protein levels in ob/+ myotubes (Fig. 6A, left panel). In myotubes isolated from ob/ob mice, SREBP-1c and FAS protein levels were very high and insulin had no additive effect (right panel). In contrast, BIO drastically downregulated SREBP-1c and FAS protein levels in ob/+ as well as in ob/ob mice myotubes. Furthermore insulin was unable to up-regulate SREBP-1c and FAS protein levels in the presence of BIO (Fig. 6A).

**Fig. 6.**
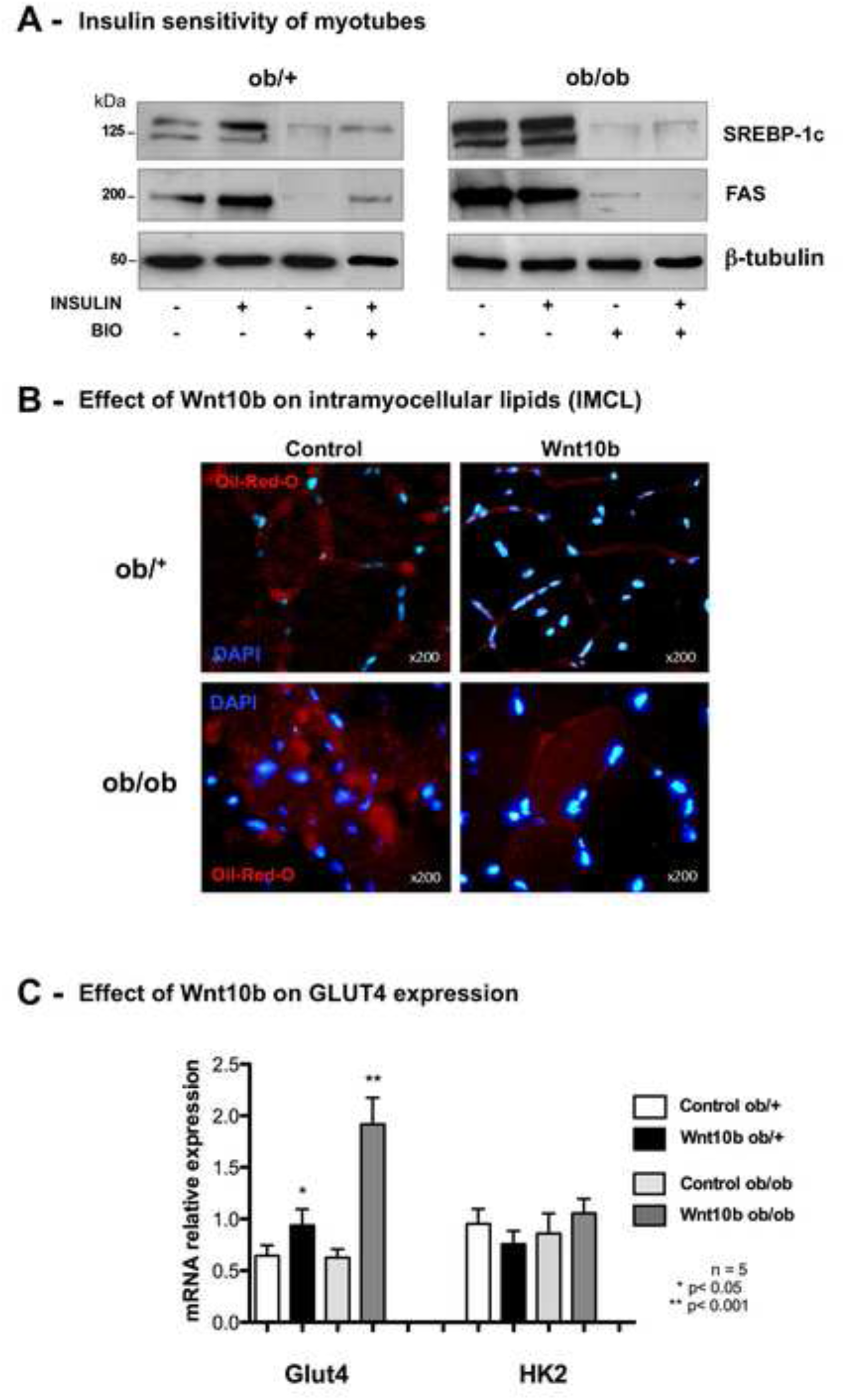
Effects of Wnt10b on glucose and lipid metabolism. **(A)** Western blot showing the effects of Wnt signaling on insulin sensitivity. Myotubes isolated from ob/+ and ob/ob mice were treated with 100 μM BIO for 24h in the presence or absence of 10 nM insulin for the last 12h. Insulin increased the lipogenic SREBP-1c and FAS protein levels in ob/+ myotubes (left panel). On the contrary myotubes from ob/ob mice were not sensitive to insulin (right panel). BIO addition drastically downregulated SREBP-1c and FAS protein levels even in the presence of insulin in ob/+ as well as in ob/ob mice myotubes. **(B)** Effect of Wnt10b on intramyocellular lipids in TA muscles. In vivo electrotransfection of Wnt10b decreased intramyocellular lipid accumulation (Oil-Red-O staining) 3 weeks after treatment, especially in ob/ob mice. Nuclei were counterstained in blue with DAPI. **(C)** Effect of Wnt10b electrotransfection on GLUT4 expression. Q-PCR quantification of *GLUT4* and *HK2* mRNA expression in TA muscles 3 weeks after Wnt10b electrotransfection. *GLUT4* mRNA was increased by 30% in ob/+ muscle whereas *GLUT4* was up-regulated by 3-fold in ob/ob muscle. In contrast, *HK2* mRNA was not affected by Wnt10b overexpression.

#### Effect of Wnt10b on intramyocellular lipid (IMCL) accumulation in TA muscles

A negative correlation between insulin sensitivity and intramyocellular lipid content has been observed in ob/ob mice [39]. To assess the role of Wnt signaling on IMCL accumulation, sections of TA muscles electrotransfected with Wnt10b were stained with Oil red O. Three weeks after transfection, Wnt10b drastically decreased IMCL accumulation, particularly in the TA of ob/ob mice (Fig. 6B).

#### Effect of Wnt10b electrotransfection on GLUT4 glucose transporter

We reported here that hyperglycemia was significantly decreased in ob/ob mice after activation of the Wnt/β-catenin pathway in TA muscles. In fact, whatever the method we used (Wnt10b overexpression or BIO injections in TA muscles), hyperglycemia was lowered by roughly 30% in the 19 ob/ob mice that were treated (Fig. 2A, 2B). In skeletal muscle, insulin stimulates glucose transport by inducing the translocation of intracellular GLUT4 glucose transporters to the plasma membrane, greatly increasing the rate of glucose transport [40]. To test whether hyperglycemia of ob/ob mice could be lowered through an increase of glucose transport, we quantified by Real-Time-PCR *GLUT4* mRNA expression in TA muscles three weeks after Wnt10b electrotransfection. *GLUT4* mRNA was up-regulated by 3-fold in the TA of ob/ob mice whereas it was only increased by 30% in the TA of ob/+ mice. In contrast, hexokinase 2 mRNA (*HK2)* which is insuline-responsive in skeletal muscle of non-diabetic subjects was not affected by Wnt10b overexpression in control and obese mice (Fig. 6C). These results suggest that Wnt10b could increase glucose transport in ob/ob mice muscle by up-regulating GLUT4 independently of insulin.

### WNT SIGNALING INCREASED THE EXPRESSION OF CYTOKINES IN MUSCLE

We hypothesized that TA muscle might secrete several factors that could exert specific endocrine effects on other organs in response to the activation of Wnt signaling.

### Wnt signaling increased interleukins expression in the TA muscle

We quantified by Real-Time PCR the expression of 3 cytokines, namely IL-4, IL-6 and IL-10 in TA muscles. BIO and Wnt10b increased by 2-to 3-fold IL-4 and IL-10 mRNA expression in TA muscles of ob/ob and ob/+ mice respectively. Similarly, IL-6 mRNA was increased by 3-to 5-fold by BIO and Wnt10b in the TA of ob/ob and ob/+ mice (Fig. 7-A).

**Fig. 7.**
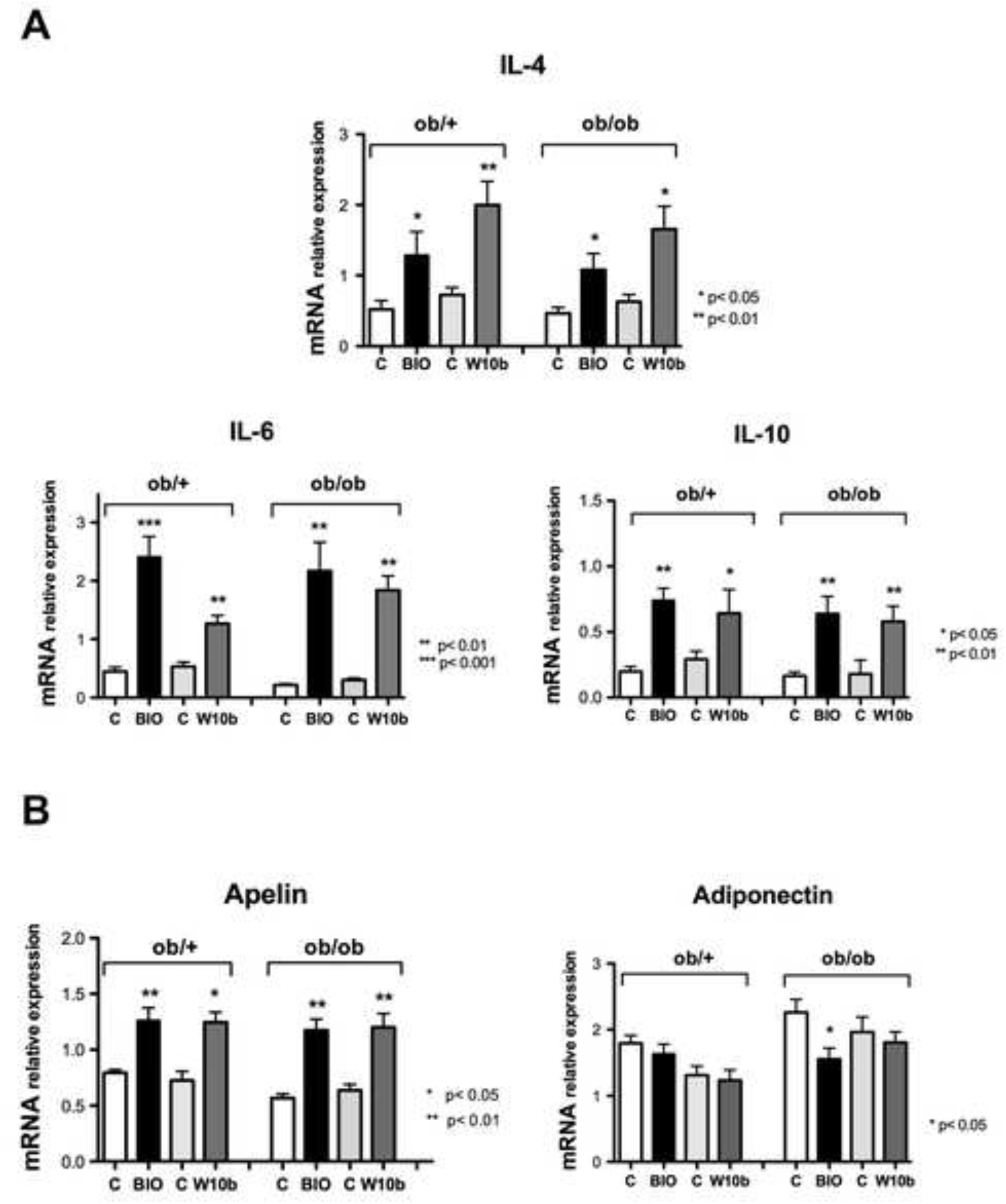
Wnt signaling increased cytokins expression in TA muscles. **(A)** Real-time PCR expression of IL-4, IL-6 and IL-10 three weeks after in vivo electrotransfection of Wnt10b or injection of BIO in TA muscles of ob/+ and ob/ob mice. BIO and Wnt10b increased by 2- and 3-fold IL-4 and IL-10 mRNA expression in TA muscles of ob/ob and ob/+ mice respectively, whereas IL-6 mRNA was increased by 3-to 5-fold. **(B)** Wnt signaling increased by 2-fold apelin mRNA expression both in ob/+ and ob/ob mice muscles. In contrast, no noteworthy difference was observed in the expression of adiponectin in TA muscles of both mouse models.

### Wnt signaling increased apelin but not adiponectin in the TA muscle

Another relevant hypothesis to explain the decrease of glycemia after activation of Wnt signaling is a role for apelin, one of the natural ligand of the orphan seven-transmembrane receptor APJ. In fact, Wnt signaling increased by 2-fold apelin mRNA expression both in ob/+ and ob/ob mice muscle. In contrast, no noteworthy difference was observed in the expression of adiponectin (a protein that modulates glucose and fatty acid regulation) in TA muscles of both mouse models (Fig. 7-B). These results strongly suggest that activation of the Wnt/μ-catenin pathway in adult myofibers induced the release of numerous factors by skeletal muscle that could play a role in glucose homeostasis.

## DISCUSSION

In this article we highlighted an important role for Wnt/μ-catenin signaling in decreasing the lipogenic factor *Srebp-1c* that we have shown to be responsible for de novo lipogenesis and insulin resistance in skeletal muscle [18, 19] along with a shift towards oxidative MHC-expressing fibers in obese and diabetic ob/ob mice. Moreover, we showed that activation of Wnt signaling in skeletal muscle led to the diminution of hyperglycemia in ob/ob mice.

### Wnt signaling induced a muscle fiber switch towards a more oxidative metabolism

Lin et al, [41] identified the peroxisome-proliferator-activated receptor-gamma co-activator PGC-1α, which is expressed in several tissues including brown fat and skeletal muscle, as an activator of oxidative metabolism. Physiological levels of PGC-1α in transgenic mice induced a fiber type conversion, as muscles normally rich in type II fibers were redder and activated genes of the mitochondrial oxidative metabolism [41]. Using fiber-type-specific promoters, these authors showed in cultured muscle cells that PGC-1α was able to activate transcription in cooperation with Mef2 protein and serves as a target for calcineurin signaling, which has been involved in slow fiber gene expression [42]. On the other hand muscle insulin resistance can coexist with mitochondrial abnormalities such as changes in morphology, reduced activity of NADH/O_2_ oxidoreductase, decreased ATP synthesis and flow of Krebs cycle, as well as lower expression of genes involved in mitochondrial biogenesis and oxidative metabolism (*Pgc-1α, Nrf -1*…) [43, 44, 45, 46]. We found that activation of Wnt signaling in TA muscle using Wnt10b overexpression or BIO injection induced a switch in fiber-type and metabolism. Oxidative fiber number (expressing MHC-I and MHC-IIa) was increased both in ob/+ and ob/ob mice muscles, while glycolytic fiber number was decreased of the same amount (Fig. 3-4). This switch was concomitant with the up-regulation of *Pgc-1α* and its targets *Hif-2α* and *Mef-2c* (Fig. 5A), all factors that are involved in mitochondrial functions [37, 47]. In contrast, *Pgc-1β* expression was unchanged (Fig. 5A). This is in accordance with what Ramamoorty et al [48] reported recently in PGC-1β^(i)skm−/−^ mice where myofiber myosin heavy chain composition and mitochondrial number, muscle strength and glucose homeostasis were unaffected. In addition, mitochondrial protein content was up-regulated (porin, subunits belonging to complex-I and -III of the respiratory chain) (Fig. 5B). Moreover, Wnt signaling increased the mitochondrial respiratory reserve capacity of muscle cells isolated from ob/+ and ob/ob mice (Fig. 5C). Rowe et al [49] showed that mice lacking both PGC-1α and -β in myocytes have profoundly deficient mitochondrial respiration but have preserved mitochondrial content and fiber-type composition. In contrast, these mice have an impaired forced exercise capacity [49]. In fact it seems that PGC-1α controls mitochondrial activity and fiber typing in response to external stimuli [50]. Our results show that Wnt10b overexpression in TA muscle mimics the positive effects of exercise on oxidative fiber number and mitochondrial respiratory reserve capacity probably through the increase of PGC1α expression.

### Wnt signaling decreased lipogenic factors and intramyocellular lipids

Activation of Wnt signaling in myofibers down-regulated lipogenic factors (SREBP-1c, FAS) and intramyocellular lipids (Fig. 6) along with a reduction in fiber CSA and an increase in oxidative metabolism, especially in ob/ob mice muscles. This result also suggests that Wnt signaling mimics endurance training. Recently, Shan et al [51] reported a role for Lkb1 (a primary upstream kinase of AMPK that is required for maintaining energy homeostasis) in diminishing ectopic lipid deposition in muscle satellite cells and their descendent mature muscles through the AMPK pathway. In fact we have previously shown that intramyocellular lipid deposition was diminished in response to the activation of Wnt signaling in mouse myotubes through a differential activation of the AMPK pathway, thus increasing insulin sensitivity and glucose uptake in insulino-resistant mouse myotubes [19]. Taken together, these results suggest that drugs that activate Wnt canonical signaling could improve insulin sensitivity in diabetic subjects.

### Wnt signaling decreased Myostatin and increased Follistatin expression in TA Muscle

Local or systemic activation of Wnt signaling through Wnt10b overexpression or BIO injection induced the down-regulation of *Mstn* mRNA expression in both ob/+ and ob/ob TA muscles, which fits what was observed in the zebrafish [52]. In contrast, *Fst* expression, a TGF-β antagonist, was only enhanced in the TA of ob/ob mice (Fig. 3). Myostatin, a negative regulator of muscle mass [35] determines both muscle fiber number, metabolism and size, and has also a regulatory role in skeletal muscle fibrosis [53]. It was shown that gene therapy using follistatin to inhibit myostatin increased muscle size and strength with reduced fibrosis in the mdx mouse, while increased fibrosis was observed in mdx mice by activation of canonical Wnt signaling through TGF-β2 activation [54] [55]. In our 3-week experiments, the decrease of myostatin expression didn’t induce a significant gain of weight in TA muscle, but showed a trend to the increase in ob/ob mice muscles (Additional File.1). In contrast, muscles of Myostatin -/- mice showed decreased aerobic and increased anaerobic metabolism [56] at the opposite of what we reported here.

These results suggest that our experiments could have been not long enough to induce an effect on muscle weight, or that the decrease in Myostatin expression and the increase in Follistatin expression could be somehow counteracted in the context of activated canonical Wnt signaling, as it was shown by Kuroda et al [57].

### Stimulation of Wnt signaling in TA muscle decreased hyperglycemia of ob/ob mice

The more striking result that we reported here was the decrease of hyperglycemia in ob/ob mice subsequently to a local activation of the Wnt/β-catenin pathway. In fact, hyperglycemia was lowered by roughly 30% in the 19 ob/ob mice we have treated. The question is : how a local stimulation of Wnt signaling could induce a systemic effect i.e. the decrease of hyperglycemia? We hypothesized that Wnt signaling could increase glucose transport in skeletal muscle of ob/ob mice that are resistant to insulin.

#### Effect of Wnt signaling on glucose transport

In skeletal muscle, insulin stimulates glucose transport by inducing the translocation of intracellular GLUT4 glucose transporters to the plasma membrane [40]. In both insulin-sensitive and insulin resistant myotubes, we have shown previously that stimulation of the Wnt/β-catenin pathway increased basal glucose transport by stimulating the translocation of GLUT4 to the plasma membrane through a pathway independent of insulin, i.e. the AMPK-α1 pathway [19]. Here we show that Wnt signaling up-regulated the glucose transporter GLUT4 particularly in the TA muscle of insulin-resistant ob/ob mice, suggesting that increased glucose uptake by skeletal muscle could partly explain the diminution of hyperglycemia.

#### Effect of Wnt signaling on interleukins expression

On the other hand, contracting muscles express and secrete several hundred proteins among which cytokines and peptides classified as "myokines" [58] that may contribute to exercise-induced protection against chronic diseases such as obesity and type 2 diabetes. We show here that BIO injections promoted IL-6, IL-10 and IL-4 mRNA expression in TA muscles of ob/ob mice and their controls (Fig. 7). It is well established that muscle fibers express the myokine IL-6 which subsequently exerts its effects both locally within the muscle to increase glucose uptake and fat oxidation [59–61] and, when released into the circulation, peripherally in several organs in a hormone-like fashion. Acute treatment of muscle cells with IL-6 increased both basal glucose uptake and translocation of the glucose transporter GLUT4 from intracellular compartments to the plasma membrane by activation of AMPK [61]. These results are in accordance with what we have previously shown in muscle cells where BIO, like IL-6, was able to increase glucose transport via GLUT4 translocation to the plasma membrane through the activation of AMPK-α1 signaling [19]. Less is known about the role of IL-4 in muscle metabolism and glucose uptake. In addition, IL-10 is an anti-inflammatory cytokine reported to prevent muscle insulin resistance by attenuating an obesity-associated macrophage and cytokine response in skeletal muscle [62]. This suggests that injections of BIO in TA muscle or intraperitoneally resulted in the up-regulation of Wnt10b and activation of Wnt signaling in the TA muscle, which could in turn induce interleukins secretion and release in the blood flow to increase glucose uptake by peripheral tissues, then down-regulating hyperglycemia. As work from several groups [63, 64] has demonstrated that IL-6 exhibits many "leptin-like" actions such as activating AMPK and insulin signaling in skeletal muscle [65], BIO could have a "leptin-like" effect by inducing interleukins secretion in skeletal muscle of ob/ob mice that are devoid of leptin. In this context our results suggest that activation of Wnt signaling through Wnt10b overexpression or BIO injections induced secretion of IL-6 along with IL-4 and IL-10 in skeletal muscle. Their release in the blood flow could increase glucose uptake by peripheral tissues promoting the decrease of hyperglycemia that we observed in ob/ob mice.

#### Wnt signaling could diminish hyperglycemia through the up-regulation of Apelin

Apelin is a circulating peptide expressed in different tissues among which skeletal muscle, but also produced and secreted by human and mouse adipocytes [66]. Recently apelin was identified as a novel myokine in obese men, where a two-fold increase in apelin mRNA level was found in muscle (but not in adipose tissue) after a 8-week endurance training program [67]. Importantly, changes in muscle apelin mRNA levels were positively related to whole-body insulin sensitivity improvement [68]. Furthermore, in obese and insulin-resistant db/db mice, apelin restored glucose tolerance and increased glucose utilization [69]. We observed a 2-fold increase in apelin mRNA expression in the TA muscle of ob/+ and ob/ob mice (Figure 7). This result suggests that BIO and Wnt10b overexpression could induce an increase in apelin secretion that might explain the down-regulation of hyperglycemia we have reported here in ob/ob mice. Nevertheless, although our results are very similar to the apelin effects mentioned above, further investigations should be performed to confirm our hypothesis.

In conclusion, activation of Wnt signaling in skeletal muscle using drugs such as BIO mimics the effect of endurance training on muscle fiber metabolism, intramyocellular lipid accumulation, release of myokines by skeletal muscle to improve glucose homeostasis in obese and diabetic ob/ob mice. As a consequence the use of GSK-3β-specific inhibitors such as BIO-derivatives that stimulate the Wnt/β-catenin pathway could provide a new insight into the treatment of obesity and/or type 2 diabetes as they normalize glycemia in hyperglycemic mice without inducing hypoglycemia in normoglycemic mice.

## AKNOWLEDGMENTS

We thank Dr. Bernardi H. (INRA, Montpellier, France), Dr. Dugail (Institut des Cordeliers, Paris-France), Dr. Lombes A. (INSERM U1016, Paris, France) and Dr. Schiaffino S. (Padova-Italy) for providing pcDNA3.1-Wnt10b plasmid, FAS antibody, complexes I/III antibodies and MHC-IIa antibody respectively. This work was supported by the « Association Française contre les Myopathies » (AFM), the Institut National de la Santé et de la Recherche Médicale (INSERM) and the Agence Nationale pour la Recherche (ANR RPV09108KKA). Eleni Christodoulou was the recipient of a research fellowship from "Région Ile-de-France ". We thank Dt Catherine Postric for helpful discussion.

---

**Additional File1:**
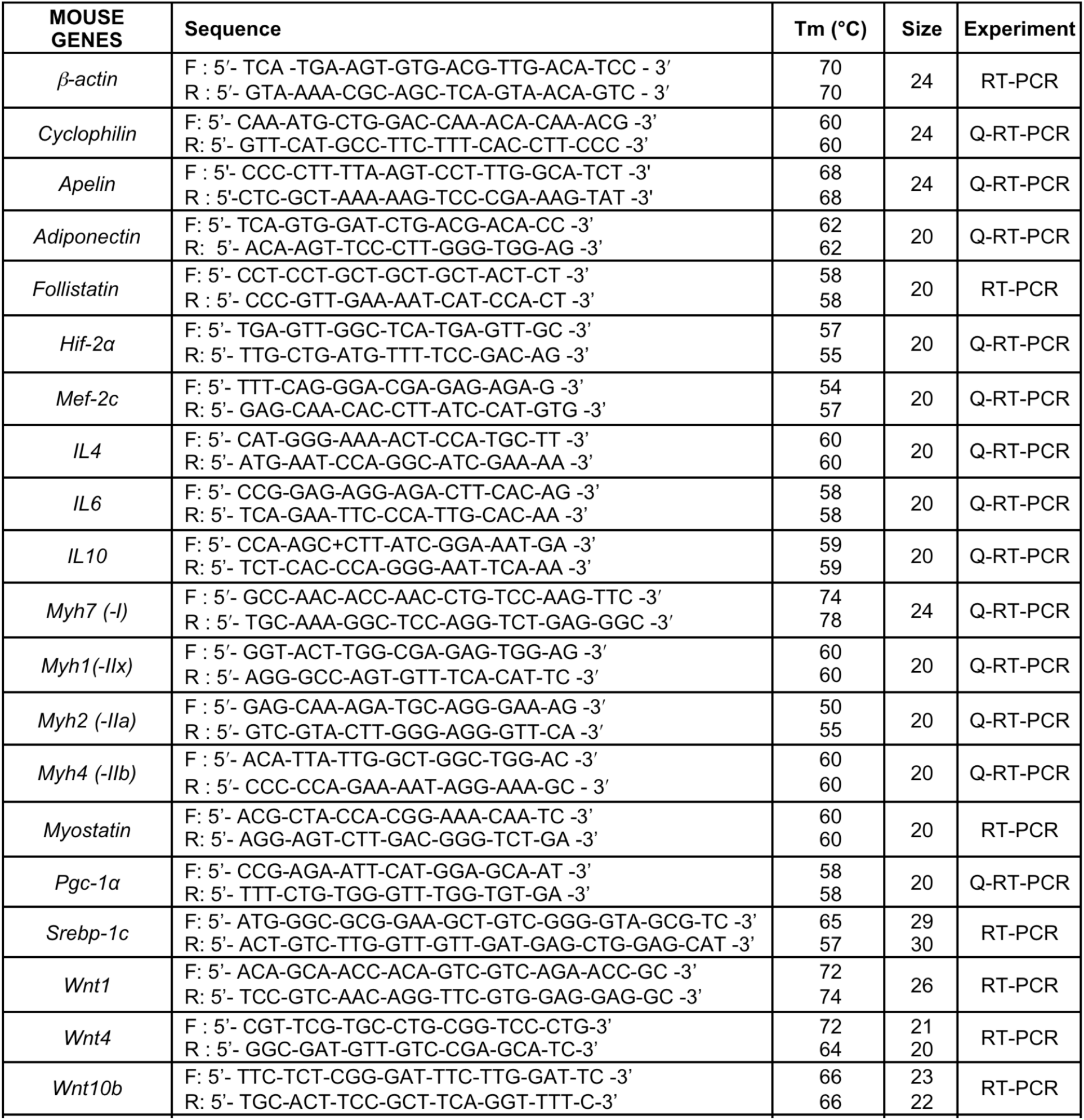
Oligonucleotides used for PCR and Q-PCR experimentss.

## REFERENCES

1. van Amerongen R, Nusse R: Towards an integrated view of Wnt signaling in development. Development 2009, 136(19):3205–3214.

2. Logan CY, Nusse R: The Wnt signaling pathway in development and disease. Annu Rev Cell Dev Biol 2004, 20:781–810.

3. Reya T, Clevers H: Wnt signalling in stem cells and cancer. Nature 2005, 434(7035):843–850.

4. Christodoulides C, Scarda A, Granzotto M, Milan G, Dalla Nora E, Keogh J, De Pergola G, Stirling H, Pannacciulli N, Sethi JK et al: WNT10B mutations in human obesity. Diabetologia 2006, 49(4):678–684.

5. Welters HJ, Kulkarni RN: Wnt signaling: relevance to beta-cell biology and diabetes. Trends Endocrinol Metab 2008, 19(10):349–355.

6. Kanazawa A, Tsukada S, Sekine A, Tsunoda T, Takahashi A, Kashiwagi A, Tanaka Y, Babazono T, Matsuda M, Kaku K et al: Association of the gene encoding wingless-type mammary tumor virus integration-site family member 5B (WNT5B) with type 2 diabetes. Am J Hum Genet 2004, 75(5):832–843.

7. Anagnostou SH, Shepherd PR: Glucose induces an autocrine activation of the Wnt/beta-catenin pathway in macrophage cell lines. Biochem J 2008, 416(2):211–218.

8. Aslanidi G, Kroutov V, Philipsberg G, Lamb K, Campbell-Thompson M, Walter GA, Kurenov S, Ignacio Aguirre J, Keller P, Hankenson K et al: Ectopic expression of Wnt10b decreases adiposity and improves glucose homeostasis in obese rats. Am J Physiol Endocrinol Metab 2007, 293(3):E726–736.

9. Longo KA, Wright WS, Kang S, Gerin I, Chiang SH, Lucas PC, Opp MR, MacDougald OA: Wnt10b inhibits development of white and brown adipose tissues. J Biol Chem 2004, 279(34):35503–35509.

10. Wright WS, Longo KA, Dolinsky VW, Gerin I, Kang S, Bennett CN, Chiang SH, Prestwich TC, Gress C, Burant CF et al: Wnt10b inhibits obesity in ob/ob and agouti mice. Diabetes 2007, 56(2):295–303.

11. Koza RA, Nikonova L, Hogan J, Rim JS, Mendoza T, Faulk C, Skaf J, Kozak LP: Changes in gene expression foreshadow diet-induced obesity in genetically identical mice. PLoS Genet 2006, 2(5):e81.

12. Ross SE, Hemati N, Longo KA, Bennett CN, Lucas PC, Erickson RL, MacDougald OA: Inhibition of adipogenesis by Wnt signaling. Science 2000, 289(5481):950–953.

13. Cossu G, Borello U: Wnt signaling and the activation of myogenesis in mammals. Embo J 1999, 18(24):6867–6872.

14. Tajbakhsh S, Borello U, Vivarelli E, Kelly R, Papkoff J, Duprez D, Buckingham M, Cossu G: Differential activation of Myf5 and MyoD by different Wnts in explants of mouse paraxial mesoderm and the later activation of myogenesis in the absence of Myf5. Development 1998, 125(21):4155–4162.

15. Virkamaki A, Korsheninnikova E, Seppala-Lindroos A, Vehkavaara S, Goto T, Halavaara J, Hakkinen AM, Yki-Jarvinen H: Intramyocellular lipid is associated with resistance to in vivo insulin actions on glucose uptake, antilipolysis, and early insulin signaling pathways in human skeletal muscle. Diabetes 2001, 50(10):2337–2343.

16. Machann J, Haring H, Schick F, Stumvoll M: Intramyocellular lipids and insulin resistance. Diabetes Obes Metab 2004, 6(4):239–248.

17. Lee JH, Bassel-Duby R, Olson EN: Heart- and muscle-derived signaling system dependent on MED13 and Wingless controls obesity in Drosophila. Proc Natl Acad Sci U S A 2014, 111(26):9491–9496.

18. Guillet-Deniau I, Pichard AL, Kone A, Esnous C, Nieruchalski M, Girard J, Prip-Buus C: Glucose induces de novo lipogenesis in rat muscle satellite cells through a sterol-regulatory-element-binding-protein-1c-dependent pathway. J Cell Sci 2004, 117(Pt 10):1937–1944.

19. Abiola M, Favier M, Christodoulou-Vafeiadou E, Pichard AL, Martelly I, Guillet-Deniau I: Activation of Wnt/beta-catenin signaling increases insulin sensitivity through a reciprocal regulation of Wnt10b and SREBP-1c in skeletal muscle cells. PloS one 2009, 4(12):e8509.

20. Shimomura I, Bashmakov Y, Ikemoto S, Horton JD, Brown MS, Goldstein JL: Insulin selectively increases SREBP-1c mRNA in the livers of rats with streptozotocin-induced diabetes. Proc Natl Acad Sci U S A 1999, 96(24):13656–13661.

21. Foretz M, Guichard C, Ferre P, Foufelle F: Sterol regulatory element binding protein-1c is a major mediator of insulin action on the hepatic expression of glucokinase and lipogenesis-related genes [see comments]. Proc Natl Acad Sci U S A 1999, 96(22):12737–12742.

22. Ducluzeau PH, Perretti N, Laville M, Andreelli F, Vega N, Riou JP, Vidal H: Regulation by insulin of gene expression in human skeletal muscle and adipose tissue. Evidence for specific defects in type 2 diabetes. Diabetes 2001, 50(5):1134–1142.

23. Guillet-Deniau I, Mieulet V, Le Lay S, Achouri Y, Carre D, Girard J, Foufelle F, Ferre P: Sterol regulatory element binding protein-1c expression and action in rat muscles: insulin-like effects on the control of glycolytic and lipogenic enzymes and UCP3 gene expression. Diabetes 2002, 51(6):1722–1728.

24. Simoneau JA, Kelley DE: Altered glycolytic and oxidative capacities of skeletal muscle contribute to insulin resistance in NIDDM. J Appl Physiol 1997, 83(1):166–171.

25. Coggan AR, Abduljalil AM, Swanson SC, Earle MS, Farris JW, Mendenhall LA, Robitaille PM: Muscle metabolism during exercise in young and older untrained and endurance-trained men. J Appl Physiol 1993, 75(5):2125–2133.

26. Papa S: Mitochondrial oxidative phosphorylation changes in the life span. Molecular aspects and physiopathological implications. Biochim Biophys Acta 1996, 1276(2):87–105.

27. Anakwe K, Robson L, Hadley J, Buxton P, Church V, Allen S, Hartmann C, Harfe B, Nohno T, Brown AM et al: Wnt signalling regulates myogenic differentiation in the developing avian wing. Development 2003, 130(15):3503–3514.

28. Hutcheson DA, Zhao J, Merrell A, Haldar M, Kardon G: Embryonic and fetal limb myogenic cells are derived from developmentally distinct progenitors and have different requirements for beta-catenin. Genes Dev 2009, 23(8):997–1013.

29. Grifone R, Laclef C, Spitz F, Lopez S, Demignon J, Guidotti JE, Kawakami K, Xu PX, Kelly R, Petrof BJ et al: Six1 and Eya1 expression can reprogram adult muscle from the slow-twitch phenotype into the fast-twitch phenotype. Mol Cell Biol 2004, 24(14):6253–6267.

30. Meijer L, Skaltsounis AL, Magiatis P, Polychronopoulos P, Knockaert M, Leost M, Ryan XP, Vonica CA, Brivanlou A, Dajani R et al: GSK-3-selective inhibitors derived from Tyrian purple indirubins. Chem Biol 2003, 10(12):1255–1266.

31. Guillet-Deniau I, Leturque A, Girard J: Expression and cellular localization of glucose transporters (GLUT1, GLUT3, GLUT4) during differentiation of myogenic cells isolated from rat foetuses. J Cell Sci 1994, 107 ( Pt 3):487–496.

32. Gnaiger E: Drug-Induced Mitochondrial Dysfunction John Wiley&Sons; 2008.

33. Koopman R, Schaart G, Hesselink MK: Optimisation of oil red O staining permits combination with immunofluorescence and automated quantification of lipids. Histochem Cell Biol 2001, 116(1):63–68.

34. Shimizu H, Julius MA, Giarre M, Zheng Z, Brown AM, Kitajewski J: Transformation by Wnt family proteins correlates with regulation of beta-catenin. Cell Growth Differ 1997, 8(12):1349–1358.

35. McPherron AC, Lawler AM, Lee SJ: Regulation of skeletal muscle mass in mice by a new TGF-beta superfamily member. Nature 1997, 387(6628):83–90.

36. Nakatani M, Takehara Y, Sugino H, Matsumoto M, Hashimoto O, Hasegawa Y, Murakami T, Uezumi A, Takeda S, Noji S et al: Transgenic expression of a myostatin inhibitor derived from follistatin increases skeletal muscle mass and ameliorates dystrophic pathology in mdx mice. Faseb J 2008, 22(2):477–487.

37. Rasbach KA, Gupta RK, Ruas JL, Wu J, Naseri E, Estall JL, Spiegelman BM: PGC-1alpha regulates a HIF2alpha-dependent switch in skeletal muscle fiber types. Proc Natl Acad Sci U S A, 107(50):21866–21871.

38. Belfiore F, Rabuazzo AM, Iannello S, Vasta D, Campione R: Insulin resistance in the obese hyperglycemic (ob/ob) mouse. Failure of hyperinsulinemia to activate hepatic pyruvate kinase (PK). Metabolism 1984, 33(2):104–106.

39. Akhmedov D, Berdeaux R: The effects of obesity on skeletal muscle regeneration. Front Physiol 2013, 4:371.

40. Mueckler M: Insulin resistance and the disruption of Glut4 trafficking in skeletal muscle. J Clin Invest 2001, 107(10):1211–1213.

41. Lin J, Wu H, Tarr PT, Zhang CY, Wu Z, Boss O, Michael LF, Puigserver P, Isotani E, Olson EN et al: Transcriptional co-activator PGC-1 alpha drives the formation of slow-twitch muscle fibres. Nature 2002, 418(6899):797–801.

42. Naya FJ, Mercer B, Shelton J, Richardson JA, Williams RS, Olson EN: Stimulation of slow skeletal muscle fiber gene expression by calcineurin in vivo. J Biol Chem 2000, 275(7):4545–4548.

43. Kelley DE, He J, Menshikova EV, Ritov VB: Dysfunction of mitochondria in human skeletal muscle in type 2 diabetes. Diabetes 2002, 51(10):2944–2950.

44. Patti ME, Butte AJ, Crunkhorn S, Cusi K, Berria R, Kashyap S, Miyazaki Y, Kohane I, Costello M, Saccone R et al: Coordinated reduction of genes of oxidative metabolism in humans with insulin resistance and diabetes: Potential role of PGC1 and NRF1. Proc Natl Acad Sci U S A 2003, 100(14):8466–8471.

45. Ritov VB, Menshikova EV, He J, Ferrell RE, Goodpaster BH, Kelley DE: Deficiency of subsarcolemmal mitochondria in obesity and type 2 diabetes. Diabetes 2005, 54(1):8–14.

46. Egan B, Zierath JR: Exercise metabolism and the molecular regulation of skeletal muscle adaptation. Cell metabolism, 17(2):162–184.

47. Potthoff MJ, Wu H, Arnold MA, Shelton JM, Backs J, McAnally J, Richardson JA, Bassel-Duby R, Olson EN: Histone deacetylase degradation and MEF2 activation promote the formation of slow-twitch myofibers. J Clin Invest 2007, 117(9):2459–2467.

48. Gali Ramamoorthy T, Laverny G, Schlagowski AI, Zoll J, Messaddeq N, Bornert JM, Panza S, Ferry A, Geny B, Metzger D: The transcriptional coregulator PGC-1beta controls mitochondrial function and anti-oxidant defence in skeletal muscles. Nat Commun 2015, 6:10210.

49. Rowe GC, Patten IS, Zsengeller ZK, El-Khoury R, Okutsu M, Bampoh S, Koulisis N, Farrell C, Hirshman MF, Yan Z et al: Disconnecting mitochondrial content from respiratory chain capacity in PGC-1-deficient skeletal muscle. Cell Rep, 3(5):1449–1456.

50. Sonoda J, Mehl IR, Chong LW, Nofsinger RR, Evans RM: PGC-1beta controls mitochondrial metabolism to modulate circadian activity, adaptive thermogenesis, and hepatic steatosis. Proc Natl Acad Sci U S A 2007, 104(12):5223–5228.

51. Shan T, Zhang P, Bi P, Kuang S: Lkb1 deletion promotes ectopic lipid accumulation in muscle progenitor cells and mature muscles. J Cell Physiol 2014.

52. Tee JM, van Rooijen C, Boonen R, Zivkovic D: Regulation of slow and fast muscle myofibrillogenesis by Wnt/beta-catenin and myostatin signaling. PloS one 2009, 4(6):e5880.

53. Li ZB, Kollias HD, Wagner KR: Myostatin directly regulates skeletal muscle fibrosis. J Biol Chem 2008, 283(28):19371–19378.

54. Rodino-Klapac LR, Haidet AM, Kota J, Handy C, Kaspar BK, Mendell JR: Inhibition of myostatin with emphasis on follistatin as a therapy for muscle disease. Muscle & nerve 2009, 39(3):283–296.

55. Biressi S, Miyabara EH, Gopinath SD, Carlig PM, Rando TA: A Wnt-TGFbeta2 axis induces a fibrogenic program in muscle stem cells from dystrophic mice. Science translational medicine 2014, 6(267):267ra176.

56. Mouisel E, Relizani K, Mille-Hamard L, Denis R, Hourde C, Agbulut O, Patel K, Arandel L, Morales-Gonzalez S, Vignaud A et al: Myostatin is a key mediator between energy metabolism and endurance capacity of skeletal muscle. American journal of physiology Regulatory, integrative and comparative physiology 2014, 307(4):R444–454.

57. Kuroda K, Kuang S, Taketo MM, Rudnicki MA: Canonical Wnt signaling induces BMP-4 to specify slow myofibrogenesis of fetal myoblasts. Skeletal muscle 2013, 3(1):5.

58. Pedersen BK, Febbraio MA: Muscle as an endocrine organ: focus on muscle-derived interleukin-6. Physiol Rev 2008, 88(4):1379–1406.

59. Bruce CR, Dyck DJ: Cytokine regulation of skeletal muscle fatty acid metabolism: effect of interleukin-6 and tumor necrosis factor-alpha. Am J Physiol Endocrinol Metab 2004, 287(4):E616–621.

60. Petersen AM, Pedersen BK: The anti-inflammatory effect of exercise. J Appl Physiol 2005, 98(4):1154–1162.

61. Carey AL, Steinberg GR, Macaulay SL, Thomas WG, Holmes AG, Ramm G, Prelovsek O, Hohnen-Behrens C, Watt MJ, James DE et al: Interleukin-6 increases insulin-stimulated glucose disposal in humans and glucose uptake and fatty acid oxidation in vitro via AMP-activated protein kinase. Diabetes 2006, 55(10):2688–2697.

62. Hong EG, Ko HJ, Cho YR, Kim HJ, Ma Z, Yu TY, Friedline RH, Kurt-Jones E, Finberg R, Fischer MA et al: Interleukin-10 prevents diet-induced insulin resistance by attenuating macrophage and cytokine response in skeletal muscle. Diabetes 2009, 58(11):2525–2535.

63. Minokoshi Y, Kim YB, Peroni OD, Fryer LG, Muller C, Carling D, Kahn BB: Leptin stimulates fatty-acid oxidation by activating AMP-activated protein kinase. Nature 2002, 415(6869):339–343.

64. Steinberg GR, Rush JW, Dyck DJ: AMPK expression and phosphorylation are increased in rodent muscle after chronic leptin treatment. Am J Physiol Endocrinol Metab 2003, 284(3):E648–654.

65. Steinberg GR, Watt MJ, Febbraio MA: Cytokine Regulation of AMPK signalling. Front Biosci 2009, 14:1902–1916.

66. Boucher J, Masri B, Daviaud D, Gesta S, Guigne C, Mazzucotelli A, Castan-Laurell I, Tack I, Knibiehler B, Carpene C et al: Apelin, a newly identified adipokine up-regulated by insulin and obesity. Endocrinology 2005, 146(4):1764–1771.

67. Dray C, Debard C, Jager J, Disse E, Daviaud D, Martin P, Attane C, Wanecq E, Guigne C, Bost F et al: Apelin and APJ regulation in adipose tissue and skeletal muscle of type 2 diabetic mice and humans. Am J Physiol Endocrinol Metab, 298(6):E1161–1169.

68. Besse-Patin A, Montastier E, Vinel C, Castan-Laurell I, Louche K, Dray C, Daviaud D, Mir L, Marques MA, Thalamas C et al: Effect of endurance training on skeletal muscle myokine expression in obese men: identification of apelin as a novel myokine. International journal of obesity 2014, 38(5):707–713.

69. Dray C, Knauf C, Daviaud D, Waget A, Boucher J, Buleon M, Cani PD, Attane C, Guigne C, Carpene C et al: Apelin stimulates glucose utilization in normal and obese insulin-resistant mice. Cell metabolism 2008, 8(5):437–445.

